# Bioinformatics analysis and validation of differentially expressed miRNA in the sperm of partners of patients with unexplained recurrent miscarriages

**DOI:** 10.1101/2024.04.09.588730

**Authors:** Hui Tian, Xiao-Xi Zhao

## Abstract

**Background:** Recurrent miscarriage is a common multifactorial disease, and even after systematic examination, the cause of the miscarriages cannot be determined in approximately 50% of patients. However, currently, few examinations are conducted for partners of patients with recurrent miscarriages. The contribution of spermatozoa to new individuals is not limited to half of the diploid genome. Recent studies have shown that spermatozoa add several components to oocytes, including a complex RNA population that plays an important role in paternal chromatin packaging, sperm maturation and empowerment, fertilisation, early embryogenesis, and developmental maintenance. In view of this, we constructed a sperm miRNA expression profile of partners of patients with unexplained recurrent miscarriages and explored the relationship between unexplained recurrent miscarriages and differential miRNA expression in sperm.

**Methods:** Semen samples were obtained from the partners of patients with unexplained recurrent miscarriages treated at the affiliated hospital of Inner Mongolia Medical University, as well as healthy men who underwent physical examinations between July 2021 and February 2023. The miRNA expression profiles of the sperm in the two groups were detected using high-throughput sequencing. Bioinformatics analysis of differentially expressed miRNAs was performed to predict target genes and their action pathways and verify the expression levels of paternal miRNAs in known embryos in the study and control groups through RT-PCR technology.

**Result:** A total of 83 differentially expressed miRNAs were detected between the two groups, of which 39 were significantly upregulated and 44 were significantly downregulated. The results showed that spermatic hsa-miR-34c-5p, hsa-miR-1285-3p, hsa-miR-486-3p, and hsa-miR-146b-3p may regulate the occurrence of unexplained recurrence through the Ras, PI3K-Akt, and MAPK signalling pathways in the bioinformatics analysis. The expression of hsa-miR-34c-5 in spouses of patients with unexplained miscarriage recurrence was downregulated (p < 0.001). Binary logistic regression analysis showed that low expression of hsa-miR-34c-5p was a high-risk factor for recurrent miscarriage (odds ratio [OR] = 4.344; 95% confidence interval [CI], 1.119–16.857; p < 0.05).

**Conclusion:** The occurrence of unexplained recurrent miscarriages is associated with the differential expression of key miRNAs in the sperms of patients’ spouses.

## BACKGROUND

Recurrent miscarriage refers to two or more consecutive miscarriages occurring under 20 weeks of gestation. The incidence of recurrent miscarriages is difficult to assess, affecting approximately 1–2% of couples [1]. It is a common multifactorial disease; however, to date, the cause of recurrent miscarriages remains unclear. Clinically, patients with recurrent miscarriages are usually examined for uterine factors, endocrine factors, immune factors, thrombosis tendencies, and chromosomal examination of both parents. After these tests, the cause of miscarriage remains undetermined in approximately 50% of couples [2]. Thus, a lack of a diagnosis prevents nearly half of patients with recurrent miscarriages from getting targeted treatment, which can cause trouble and confusion in patients and families.

Fifty percent of an embryo’s genetic material comes from the sperm. The genetic material of a spermatozoon is crucial in normal embryonic development, differentiation, embryo implantation, and placenta formation [3,4]. Small non-coding RNAs (sncRNAs) originate from gonadal cells [5]. Environmental factors can influence the sncRNA abundance or transport processes in gonadal cells, resulting in changes in the sncRNA payload of sperm [6,7]. At fertilisation, these altered sncRNAs are released into the ovum, which affects the offspring’s phenotype and health by controlling the expression of the embryonic genome [8,9].

At present, the main clinical examination for recurrent miscarriages consists of the evaluation of women, and little attention is paid to the influence of male factors on recurrent miscarriages. Based on the regulatory role of miRNAs in sperm during embryonic development, this study constructed a miRNA expression profile of the partners of patients with unexplained recurrent miscarriages using high-throughput sequencing technology. In addition, we predicted target genes and pathways through bioinformatics analysis and validated key differentially expressed miRNAs to elucidate the relationship between miRNA differential expression in spermatozoa and recurrent miscarriages, providing a scientific basis for diagnosing and treating patients with recurrent miscarriages.

## METHODS

### Participants

This study was approved by the Medical Ethics Committee of Inner Mongolia Medical University, Hohhot, China (Ethics Approval Number: YKD202301105). The selected participants provided informed consent. The semen samples were obtained from the partners of patients with unexplained recurrent miscarriages who were treated at the affiliated hospital of Inner Mongolia Medical University, as well as healthy men who underwent physical examinations between July 2021 and February 2023. This study included 19 partners of patients with two or more spontaneous miscarriages who were systematically screened for causes of miscarriage without significant abnormalities. The control group comprised 19 men who underwent pregnancy preparation physical examinations and who had no obvious abnormalities found on their semen routine examinations. Sexual activity was discontinued for 3-7 days before collecting specimens.

Three samples from the case and control groups were selected for high-throughput sequencing and bioinformatics analysis. The remaining specimens were used for key differential expression miRNA validation.

### Data and specimen collection

General patient information was obtained from the case registration information at the time of the patient’s visit.

Specimens were collected. Liquefaction was performed in a 36.6°C incubator for 30 min; 2 mL semen from each sample was purified by density gradient centrifugation, treated with somatic cell lysate buffer, and then frozen at −80°C. RNA was extracted from sperm samples using the miRNeasy Micro Kit (QIAGEN, Germany), and 1 μL of RNA was used for RNA quantification and purity assessment using an ultraviolet-visible range spectrophotometer.

### High-throughput sequencing and bioinformatics analysis

The qualifying RNA samples were subjected to reverse transcription, PCR amplification, purification of cDNA constructs, recovery of the purified constructs, and inspection for library quality before sequencing. Small RNA sequencing data were processed, and miRNAs were screened. The measurement index was calculated using transcript-per-million values to represent miRNA expression. The fold change of miRNAs between the two groups was analysed using DESeq2.0 software, and miRNAs with fold changes >1.5 and p-values <0.05 were selected as differentially expressed miRNAs. The miRanda bioinformatics algorithm was used to predict the target genes of differentially expressed miRNAs. Gene Ontology (GO) enrichment analysis and Kyoto Encyclopedia of Genes and Genomes (KEGG) enrichment analysis of the target genes were conducted.

### Verification by quantitative real-time PCR

RNA that passed the quality test underwent RNA polyadenylation and cDNA synthesis according to the instructions of the kit manufacturer (Mir-X™ miRNA First-Strand Synthesis Kit; Takara). The reaction system was configured according to the real-time fluorescence quantitative PCR user manual (ChamQ Universal SYBR® qPCR Master Mix; Vazyme), HOMO U6 was used as the endogenous reference, and the relative expression level of the target miRNA was calculated. The reaction conditions were as follows: preheating at 95°C for 30 s, followed by 40 cycles at 95°C for 10 s and 60°C for 30 s. Denaturation was conducted at 95°C for 15 s, 60°C for 60 s, and 95°C for 15 s. The sequence of the target miRNA was as follows:

hsa-miR-375-3p AGTTTGTTCGTTCGGCTC

hsa-miR-25-3p CATTGCACTTGTCTCGGTCTGA

hsa-miR-34c-5p AGGCAGTGTAGTTAGCTGATTGC

HOMO U6 GGAACGATACAGAGAAGATTAGC

### Statistical methods

Statistical analyses were performed using SPSS 19.0 software (SPSS Inc., IL, USA). The data representing age, sperm concentration, total sperm motility, and control groups were normally distributed, and between-group differences were assessed using the t-test. The RNA concentration, miR-34c, miR-375, and miR-25 expression quantity did not follow the normal distribution; therefore, between-group differences were assessed using the Mann–Whitney U test. Binary logistic regression was used to evaluate the relationship between miRNA differential expression in sperm and unexplained recurrent miscarriages. Statistical significance was set at p-values < 0.05.

## RESULTS

### Screening for differentially expressed miRNAs in the sperm of partners of patients with unexplained recurrent miscarriages

Table 1 shows the general information comparison between the case and control groups. Compared with the control group, the recurrent miscarriage group had 83 differentially expressed miRNAs, of which 39 were significantly upregulated and 44 were significantly downregulated (Table 2 and Fig 1). Of 83 differentially expressed miRNAs, 68 were successfully predicted to have 14,614 target genes and 17,111 target gene binding sites. High scores were obtained in the prediction analysis of target genes for hsa-miR-486-3p, hsa-miR-1285-3p, and hsa-miR-146b-3p.

**Table 1.** Basic characteristics of the URSM and control groups.

| Grouping | Identifier | Age | Sperm concentration<br>(10<sup>6</sup>/mL) | Total sperm motility (%) | DNA integrity (%) |
| --- | --- | --- | --- | --- | --- |
| Recurrent miscarriage group (s) | s-01 | 36 | 31.66 | 59.66 | 87 |
|  | s-02 | 30 | 40.12 | 37.85 | 90 |
|  | s-03 | 32 | 41.34 | 62.57 | 88 |
| Control group (d) | d-04 | 35 | 213.57 | 72.67 | 89 |
|  | d-05 | 39 | 138 | 55.12 | 86 |
|  | d-06 | 37 | 22.11 | 59.83 | 92 |

URSM, unexplained recurrent spontaneous miscarriage. Sample information for high-throughput sequencing

**Table 2.**
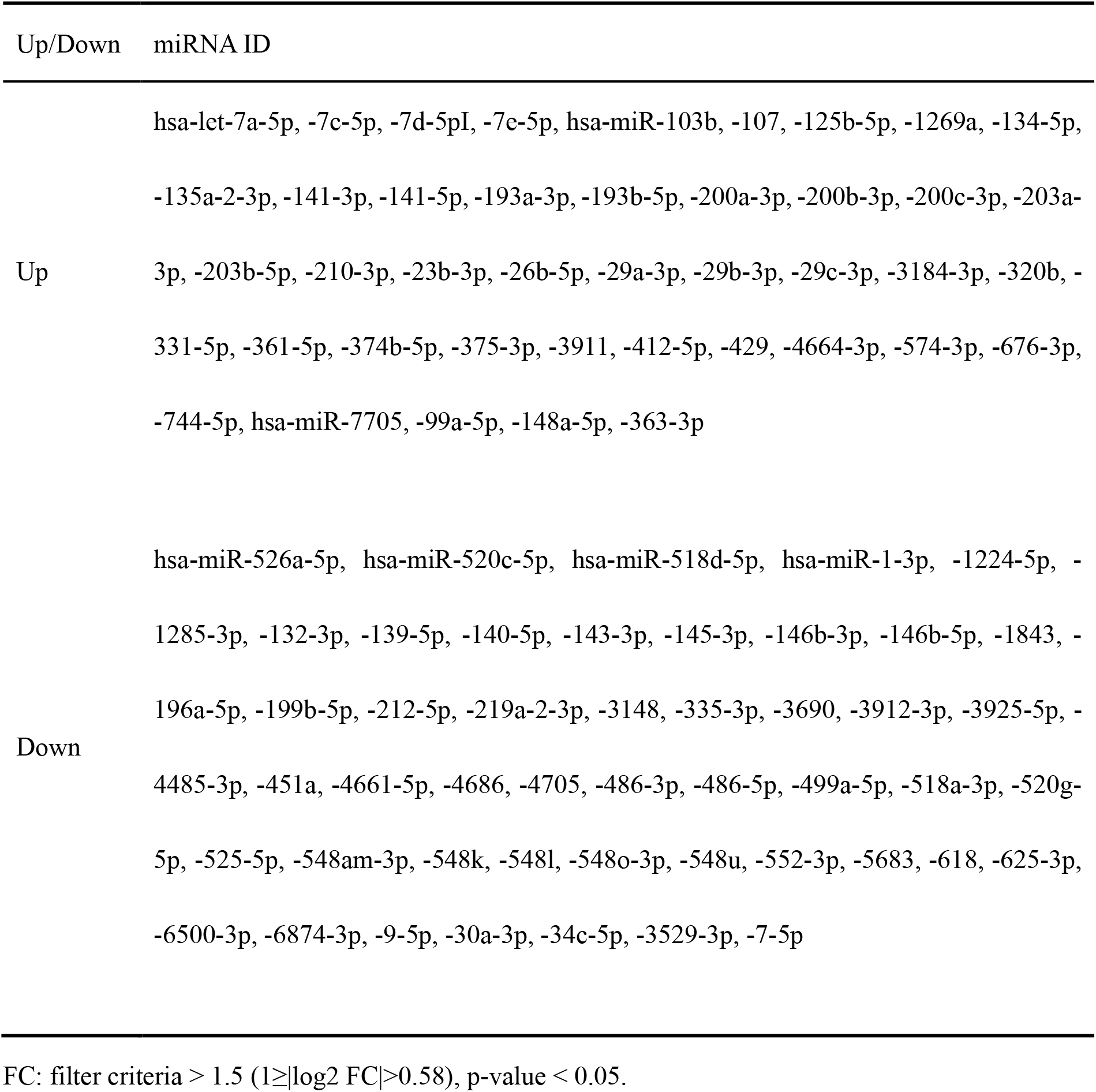
Differentially expressed miRNAs.

**Figure 1.**
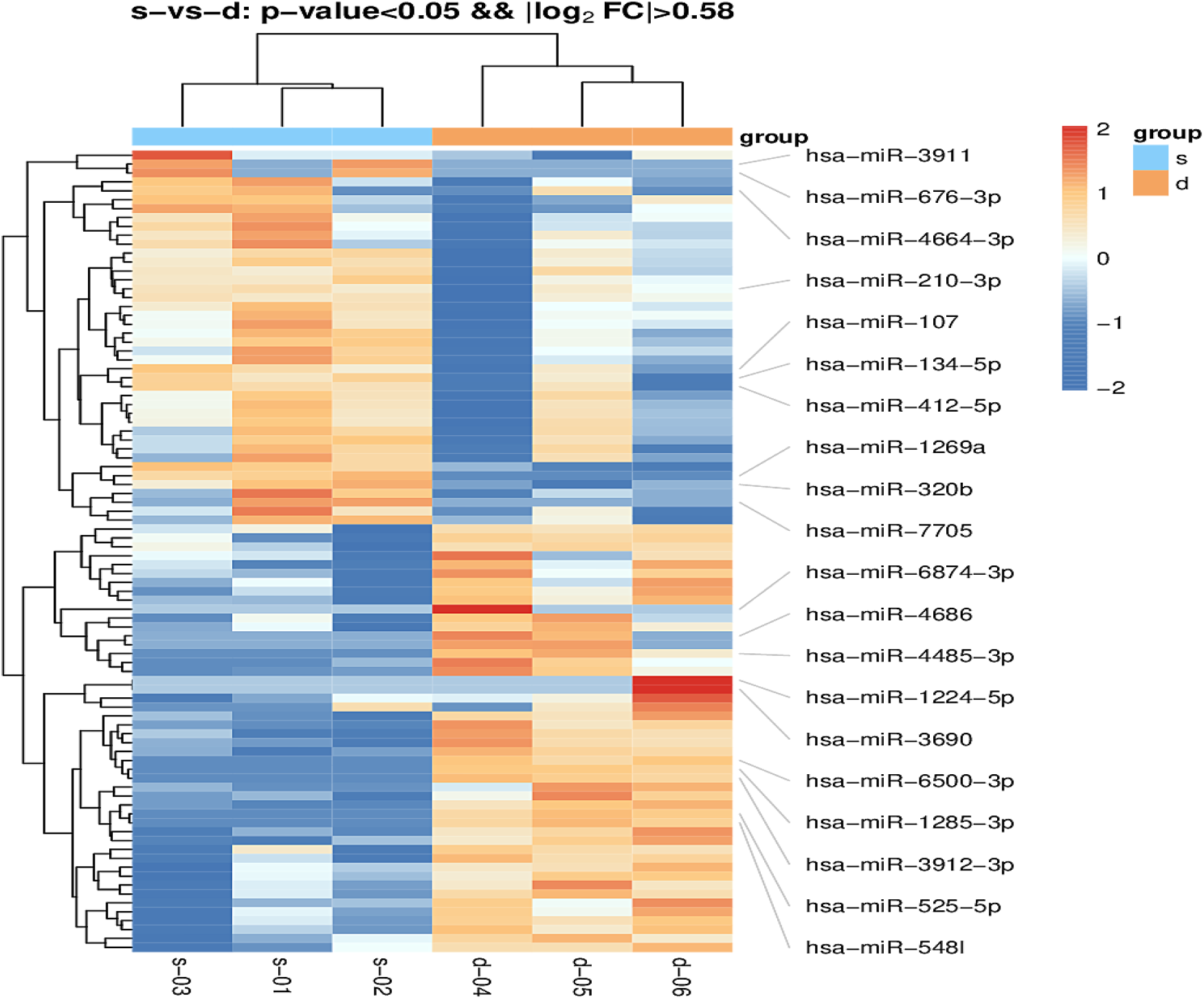
Heat map of differential miRNA expression between the URSM and control groups. Red represents upregulation, and blue represents downregulation of miRNA expression. The figure only shows the IDs of some miRNAs. URSM, unexplained recurrent spontaneous miscarriage.

### Bioinformatics analysis of differentially expressed miRNAs in sperm

The GO function enrichment analysis revealed that the differentially expressed miRNA target genes were mainly enriched in cellular processes, biological regulation, metabolic processes, multicellular organismal processes, developmental processes, cellular organelles, macromolecular complexes, membranes, binding, catalytic activity, transporter activity, and molecular transducer activity (Fig 2).

**Figure 2.**
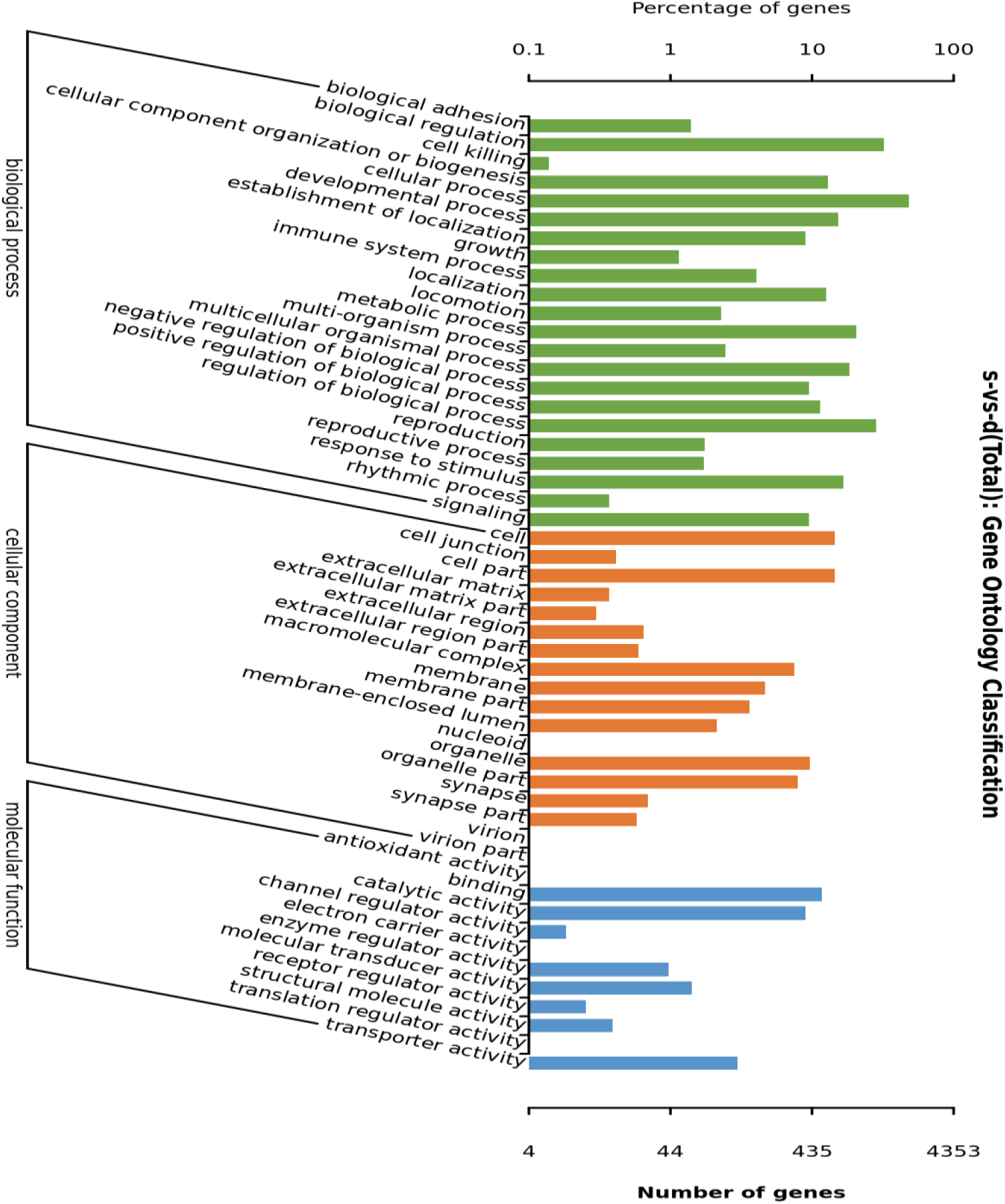
Distribution map of differentially expressed miRNA target genes and all genes at GO level 2. The vertical axis represents the name of the entry, and the horizontal axis represents the number of differentially expressed miRNA target genes for the corresponding entry. Target genes were mainly enriched in cellular processes, biological regulation, metabolic processes, multicellular organismal processes, developmental processes, cellular organelles, macromolecular complexes, membranes, binding, catalytic activity, transporter activity, and molecular transducer activity.

KEGG pathway analysis revealed that the target genes were mainly involved in signal transduction, the endocrine system, the immune system, transport and catabolism, signalling molecules and interactions, and cell growth and death (Fig 3). Among these, the target genes involved in signal transduction were the most enriched, whereas the PI3K-Akt, MAPK, and Ras signalling pathways had the most enriched target genes.

**Fig 3.**
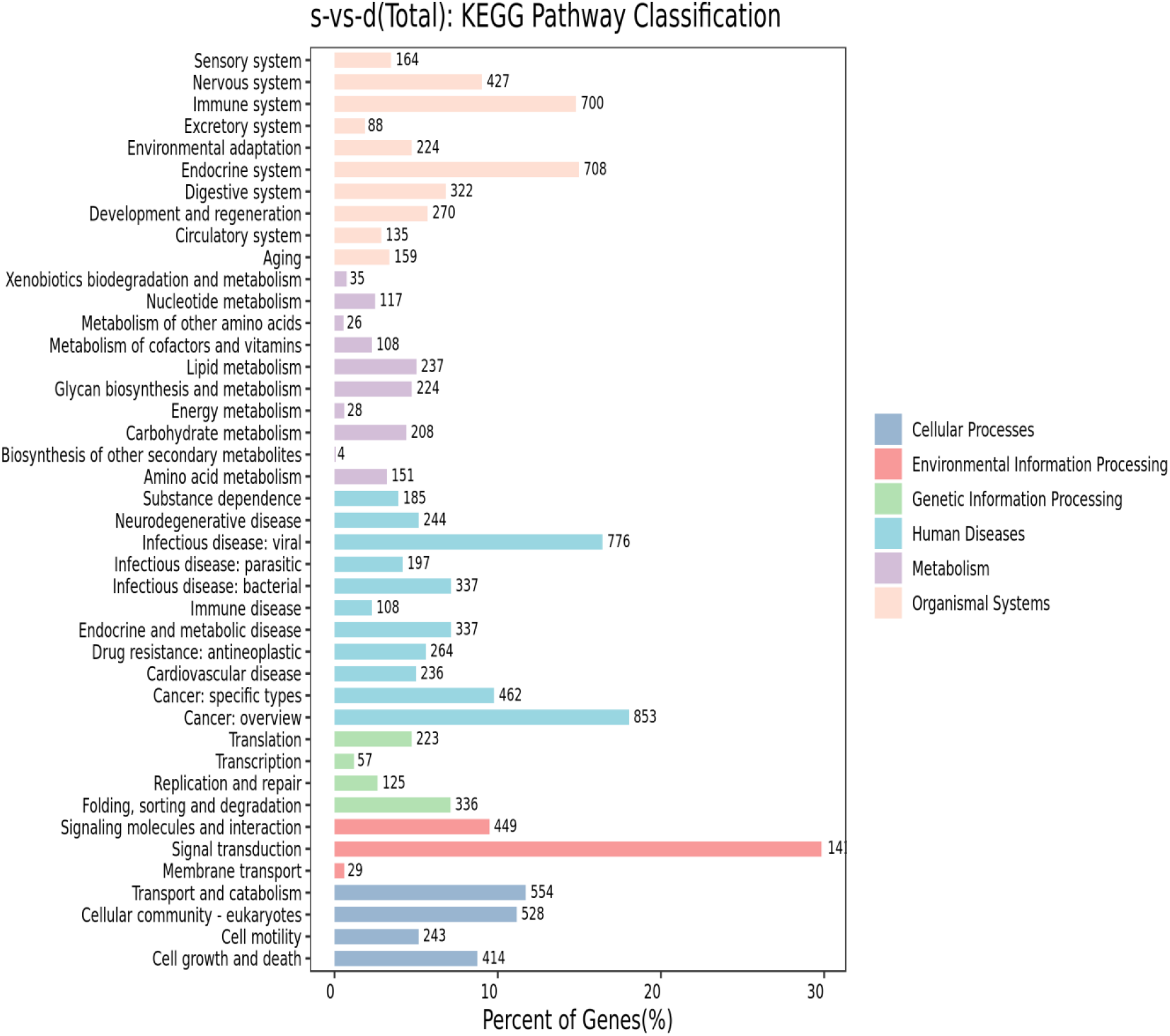
KEGG level 2 distribution map of differentially expressed miRNA target genes. The horizontal axis represents the total number and corresponding ratio of differential miRNA target genes annotated to each Level 2 pathway, the vertical axis represents the name of the Level 2 pathway, and the number represents the number of differential miRNA target genes annotated to that Level 2 pathway. Target genes are concentrated in signal transduction, the endocrine system, the immune system, transport and catabolism, signalling molecules and interactions, and cell growth and death.

### Expression of embryonic paternal miRNA in the sperm of the spouses of patients with unexplained recurrent spontaneous miscarriages (URSM)

The basic information of the case and control groups is shown in Table 3. The expression of hsa-miR-34c-5p was significantly downregulated in the recurrent miscarriage group compared to that in the control group (p < 0.01), and no significant difference was observed in hsa-miR-375-3p (p = 0.985) or hsa-miR-25-3p (p = 0.545) expression levels (Table 3).

**Table 3.**
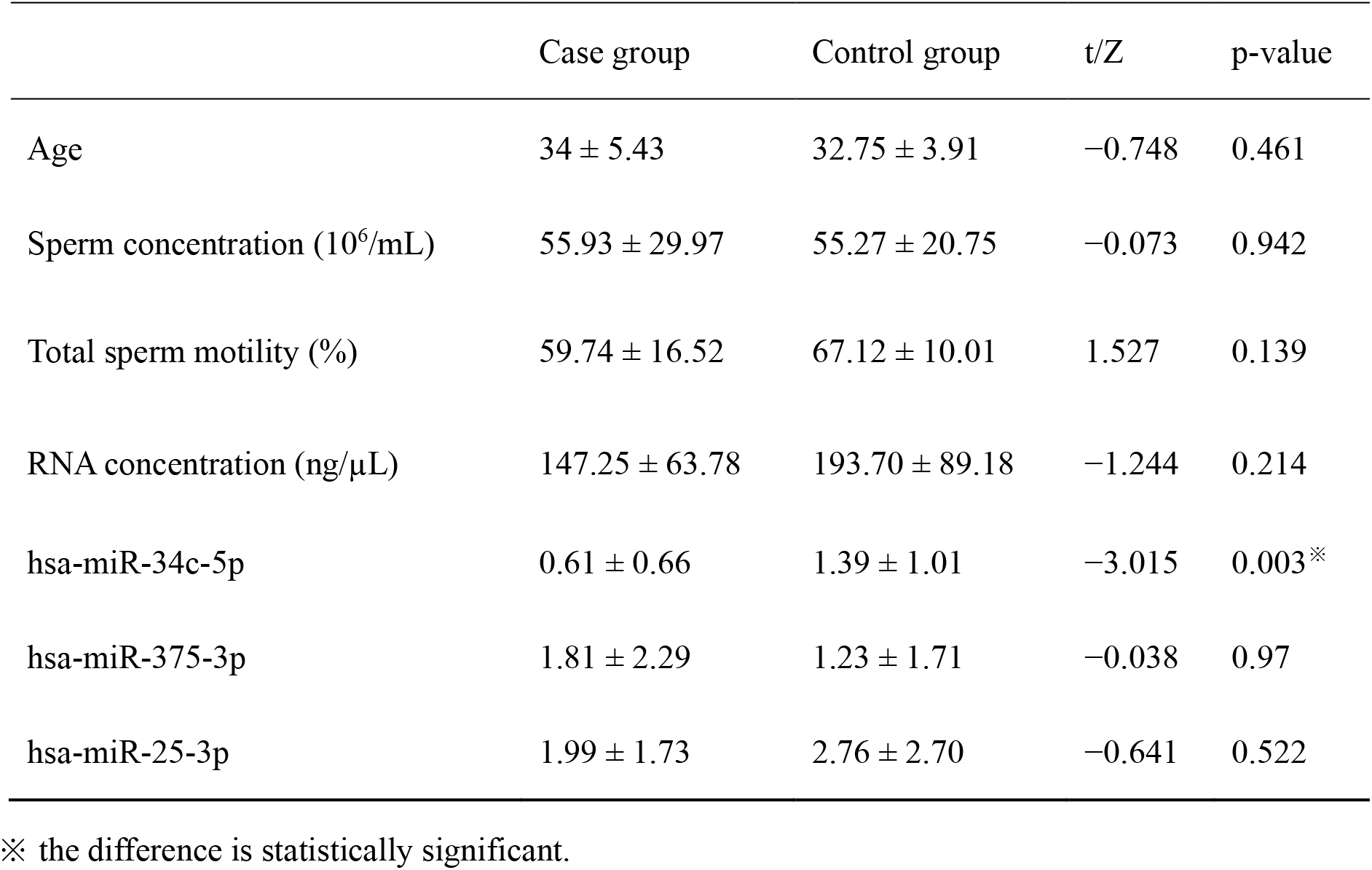
Semen parameters and expression levels of miRNAs compared between the recurrent miscarriage and control groups.

The results of the binary logistic regression analysis showed that low expression of has-miR-34c-5p is a high-risk factor for recurrent miscarriages (odds ratio [OR] = 4.344; 95% confidence interval [CI], 1.119-16.857 p < 0.05) (Table 4).

**Table 4.** Logistic regression analysis results of miR-34 expression level in sperm and risk of recurrent miscarriages.

|  |  |  |  |  | OR (95% CI) |  |  |
| --- | --- | --- | --- | --- | --- | --- | --- |
|  | B | S.E. | Wals | P-value | OR | Lower | Upper |
| miR-34 | 1.469 | 0.692 | 4.507 | 0.034<sup>※</sup> | 4.344 | 1.119 | 16.857 |
| Constant | 1.331 | 0.688 | 3.745 | 0.053 | 3.784 |  |  |

※ the difference is statistically significant.
OR: odds ratio, CI: confidence interval

## DISCUSSION

We observed differences in the expression of miRNAs between the sperm of partners of patients with URSMs and those in the control group. Bioinformatics analysis revealed that differentially expressed miRNAs are involved in some signalling pathways that may be the cause of URSMs. Previous studies have shown that other differentially expressed miRNAs in sperm play roles in embryonic development, RSM occurrence, and other processes. Examples include miR-21-5p, miR-141-3p, and miR-210-3p [10]. However, whether miRNA is differentially expressed in the spousal sperm of patients with recurrent miscarriages has not been reported.

In the target gene prediction results, hsa-miR-1285-3p, hsa-miR-486-3p, and hsa-miR-146b-3p obtained high scores. These miRNAs are expressed in the placenta, trophoblasts, and embryo [11,12,13]. Their downstream target genes, JUN, HDAC2, TP53, and FGFR3, are involved in fertilisation, early embryonic development, and other processes [14,15,16,17]. The GO functional enrichment analysis revealed that the target genes were enriched in cellular processes, biological regulation, metabolic processes, multicellular organismal and developmental processes, cellular organelles, macromolecular complexes, membranes, binding, catalytic activity, transporter activity, and molecular transducer activity. KEGG pathway enrichment analysis revealed that the pathways involving the target genes included signal transduction, endocrine system, immune system, transport and catabolism, signalling molecules and interactions, and cell growth and death. Among these, most of the target genes were involved in signal transduction, totalling 1410. The PI3K-Akt, MAPK, and Ras signalling pathways had the most enriched target genes.

The PI3K-Akt, MAPK, and Ras signalling pathways are involved in embryonic development. Ding et al. [18] discovered that circCREBBP targets miR-143-3p and promotes boar sperm survival and vitality through the PI3K-Akt signalling pathway. Zhao et al. [19] revealed that high melatonin levels may induce cell apoptosis, arrest the cell cycle, and inhibit cell growth through the PI3K-AKT signalling pathway, thus interfering with rat embryonic heart development. Nucleotide-binding oligomerisation domains 1 and 2 (NOD1 and NOD2) in the villous tissue of patients with RSMs can activate the MAPK/p38 pathway, thereby inhibiting the invasion of trophoblast cells and leading to RSM occurrence [20]. Hsa-miR-1285-3p, hsa-miR-486-3p, and hsa-miR-146b-3p target genes were also enriched in the p53 signalling pathway. Shang et al. [21] discovered that the activation of the p53/CDKN1A and p53/Bax pathways causes RSM in the villous tissues of patients. The placental villi of patients with URSMs express a high level of p53, which may result in cell apoptosis and lead to RSMs [21].

In addition, some immune-related signalling pathways, including the NOD-like receptor signalling pathway, toll-like receptor signalling pathway, B cell receptor signalling pathway, and other target genes, are enriched in the target genes of differentially expressed sperm miRNAs. Maintenance of pregnancy requires the maternal immune system to maintain tolerance of the fetus, and immune dysfunction of the maternal-fetal interface leads to recurrent miscarriages and pregnancy complications. In the nervous system, only the neurotrophin signalling pathway significantly enriches differentially expressed sperm miRNA target genes and is important in oocyte survival and developmental competence [22]. Endocrine dysfunction is an important factor in RSM occurrence. Previous studies revealed that the differentially expressed sperm miRNA target genes were enriched in the relaxin signalling pathway, thyroid hormone signalling pathway, and aldosterone synthesis and secretion pathways [23,24].

Krawetz et al. [25] discovered that hsa-miR-34c, hsa-miR-25, and hsa-miR-375 are expressed in sperm and zygotes but not in the ovary when studying human sperm miRNAs, indicating that these miRNAs are carried into the zygote by the sperm. Therefore, we selected these three miRNAs to verify their expression in the sperm of the spouses of patients with URSM.

High-throughput sequencing and RT-PCR validation confirmed the downregulation of hsa-miR-34c-5p expression in the sperm of spouses of patients with URSMs, suggesting that hsa-miR-34c is involved in early embryonic development and, thus, affects pregnancy outcomes. Many studies have confirmed the role of miR-34c in human reproduction. The expression of miR-34b and miR-34c in sperm was significantly higher in the live-birth group of patients who underwent intracytoplasmic sperm injection with teratospermia than in the non-live-birth group [26]. Yuan et al. [27] and Wu et al. [28] reported that the synergy between miR-34b/c and miR-449 is an important regulatory mechanism in spermatogenesis. The expression levels of miR-34c-5p in the sperm of men with unexplained infertility and miR-34b in the sperm of patients with oligoasthenozoospermia were significantly downregulated compared to those in the control group [29]. In the seminal plasma of patients with non-obstructive azoospermia, hsa-miR-34c-5p expression was significantly lower than that in the control group [30]. In addition, Shi et al. [31] discovered that spermatic miR-34c expression was positively correlated with the blastocyst formation rate, high-quality blastocyst rate, and pregnancy rate in a study of human embryo development during *in vitro* fertilisation. Thus, a significant amount of data has confirmed that miR-34c may be a characteristic marker of male infertility and plays a positive role in embryonic development and protection of pregnancy status.

Bioinformatics analysis revealed that the target gene of hsa-miR-34c-5p was significantly enriched in the Ras signalling pathway that is known to play a role in miscarriages and embryo implantation. Maternally Expressed Gene 3 (MEG3) inhibits RASA1 expression, activating the RAS-MAPK pathway, enhancing the proliferative and invasive abilities of trophoblasts, and facilitating miscarriages [32]. The Ras signalling pathway is regulated by Talin1 (a local adhesion complex protein necessary for cell adhesion and movement), thereby enhancing endometrial cell adhesion and promoting embryo implantation [33].

Further studies need to demonstrate the mechanism of miR-34 regulation of target genes and large sample verification and finally clarify the relationship between key differential expressions of miRNA in sperm and unexplained recurrent miscarriages.

## CONCLUSION

The occurrence of unexplained recurrent miscarriages is associated with the differential expression of key miRNAs in the sperms of patients’partners.

## Abbreviations

CI: confidence interval
FC: filter criteria
GO: Gene Ontology
KEGG: Kyoto Encyclopedia of Genes and Genomes
MEG3: maternally expressed gene
NOD1: nucleotide-binding oligomerization domain 1
NOD2: nucleotide-binding oligomerization domain 2
OR: odds ratio
RSM: recurrent spontaneous miscarriage
sncRNA: small non-coding RNA
URSM: unexplained recurrent spontaneous miscarriage

## DECLARATIONS

### Ethics approval and consent to participate

This study was approved by the Medical Ethics Committee of Inner Mongolia Medical University, Hohhot, China (Ethics Approval Number: YKD202301105). The selected participants provided informed consent.

### Consent for publication

Not applicable.

### Availability of data and materials

All data generated during this study are available at ResMan (www.medresman.org)

### Authors’ contributions

Hui Tian performed the experiments and wrote the manuscript. Xiao Xi Zhao conceived and designed the study and revised the manuscript. All the authors approved the final manuscript.

## Acknowledgements

We thank Xiujuan Chen and Yu Zhang for their support and assistance in the specimen collection.

## REFERENCES

1. ESHRE Guideline Group on RPL; Bender Atik R, Christiansen OB, Elson J, Kolte AM, Lewis S, et al. ESHRE guideline: recurrent pregnancy loss: an update in 2022. Hum Reprod Open. 2023;2023:hoad002. doi:10.1093/hropen/hoad002

2. Arias-Sosa LA, Acosta ID, Lucena-Quevedo E, Moreno-Ortiz H, Esteban-Pérez C, Forero-Castro M. Genetic and epigenetic variations associated with idiopathic recurrent pregnancy loss. J Assist Reprod Genet. 2018;35:355–66. doi:10.1007/s10815-017-1108-y

3. Pourmasumi S, Sabeti P, Ghasemi N. Male factor testing in recurrent pregnancy loss cases: A narrative review. Int J Reprod Biomed. 2022;20:447–60. doi:10.18502/ijrm.v20i6.11440

4. Giacone F, Cannarella R, Mongioì LM, Alamo A, Condorelli RA, Calogero AE, et al. Epigenetics of Male Fertility: Effects on Assisted Reproductive Techniques. World J Mens Health. 2019;37:148–56. doi:10.5534/wjmh.180071

5. Nixon B, Stanger SJ, Mihalas BP, Reilly JN, Anderson AL, Tyagi S, et al. The microRNA signature of mouse spermatozoa is substantially modified during epididymal maturation. Biol Reprod. 2015;93:91. doi:10.1095/biolreprod.115.132209

6. Chen Q, Yan W, Duan E. Epigenetic inheritance of acquired traits through sperm RNAs and sperm RNA modifications. Nat Rev Genet. 2016;17:733–43. doi:10.1038/nrg.2016.106

7. Yang C, Zeng QX, Liu JC, Yeung WS, Zhang JV, Duan YG. Role of small RNAs harbored by sperm in embryonic development and offspring phenotype. Andrology. 2022;11:770–82. doi:10.1111/andr.13347

8. Guo L, Chao SB, Xiao L, Wang ZB, Meng TG, Li YY, et al. Sperm-carried RNAs play critical roles in mouse embryonic development. Oncotarget. 2017;8:67394–405. doi:10.18632/oncotarget.18672

9. Gannon JR, Emery BR, Jenkins TG, Carrell DT. The sperm epigenome: implications for the embryo. Adv Exp Med Biol. 2014;791:53–66. doi:10.1007/978-1-4614-7783-9_4

10. Kochhar P, Dwarkanath P, Ravikumar G, Thomas A, Crasta J, Thomas T, et al. Placental expression of miR-21-5p, miR-210-3p and miR-141-3p: relation to human fetoplacental growth. Eur J Clin Nutr. 2022;76:730–8. doi:10.1038/s41430-021-01017-x

11. Xu N, Zhou X, Shi W, Ye M, Cao X, Chen S, et al. Integrative analysis of circulating microRNAs and the placental transcriptome in recurrent pregnancy loss. Front Physiol. 2022;13:893744. doi:10.3389/fphys.2022.893744

12. Zeng H, Fu Y, Shen L, Quan S. MicroRNA signatures in plasma and plasma exosome during window of implantation for implantation failure following in-vitro fertilization and embryo transfer. Reprod Biol Endocrinol. 2021;19:180. doi:10.1186/s12958-021-00855-5

13. Taga S, Hayashi M, Nunode M, Nakamura N, Ohmichi M. miR-486-5p inhibits invasion and migration of HTR8/SVneo trophoblast cells by down-regulating ARHGAP5. Placenta. 2022;123:5–11. doi:10.1016/j.placenta.2022.04.004

14. Palomares AR, Castillo-Domínguez AA, Ruiz-Galdón M, Rodriguez-Wallberg KA, Reyes-Engel A. Genetic variants in the p53 pathway influence implantation and pregnancy maintenance in IVF treatments using donor oocytes. J Assist Reprod Genet. 2021;38:3267–75. doi:10.1007/s10815-021-02324-9

15. Okumu LA, Forde N, Mamo S, McGettigan P, Mehta JP, Roche JF, et al. Temporal regulation of fibroblast growth factors and their receptors in the endometrium and conceptus during the pre-implantation period of pregnancy in cattle. Reproduction. 2014;147:825–34. doi:10.1530/REP-13-0373

16. Ma P, Schultz RM. HDAC1 and HDAC2 in mouse oocytes and preimplantation embryos: Specificity versus compensation. Cell Death Differ. 2016;23:1119–27. doi:10.1038/cdd.2016.31

17. Liu X, Zhao J, Luan X, Li S, Zhai J, Liu J, et al. SPARCL1 impedes trophoblast migration and invasion by down-regulating ERK phosphorylation and AP-1 production and altering EMT-related molecule expression. Placenta. 2020;89:33–41. doi:10.1016/j.placenta.2019.10.007

18. Ding N, Zhang Y, Huang M, Liu J, Wang C, Zhang C, et al. Circ-CREBBP inhibits sperm apoptosis via the PI3K-Akt signaling pathway by sponging miR-10384 and miR-143-3p. Commun Biol. 2022;5:1339. doi:10.1038/s42003-022-04263-2

19. Zhao A, Zhao K, Xia Y, Lyu J, Chen Y, Li S. Melatonin inhibits embryonic rat H9c2 cells growth through induction of apoptosis and cell cycle arrest via PI3K-AKT signaling pathway. Birth Defects Res. 2021;113:1171–81. doi:10.1002/bdr2.1938

20. Wang Z, Liu M, Nie X, Zhang Y, Chen Y, Zhu L, et al. NOD1 and NOD2 control the invasiveness of trophoblast cells via the MAPK/p38 signaling pathway in human first-trimester pregnancy. Placenta. 2015;36:652–60. doi:10.1016/j.placenta.2015.03.004

21. Shang W, Shu MM, Liu M, Wang AM, Lv LB, Zhao Y, et al. Elevated expressions of p53, CDKNA1, and Bax in placental villi from patients with recurrent spontaneous abortion. Eur Rev Med Pharmacol Sci. 2013;17:3376–80

22. De Sousa PA, da Silva SJ, Anderson RA. Neurotrophin signaling in oocyte survival and developmental competence: a paradigm for cellular toti-potency. Cloning Stem Cells. 2004;6:375–85. doi:10.1089/clo.2004.6.375

23. Piccirilli D, Baldini E, Massimiani M, Camaioni A, Salustri A, Bernardini R, et al. Thyroid hormone regulates protease expression and activation of Notch signaling in implantation and embryo development. J Endocrinol. 2018;236:1–12. doi:10.1530/JOE-17-0436

24. Wiegel RE, Danser AHJ, van Duijn L, Willemsen SP, Laven JSE, Steegers EAP, et al. Circulating maternal prorenin and oocyte and preimplantation embryo development: the Rotterdam Periconception Cohort. Hum Reprod. 2023;38:582–95. doi:10.1093/humrep/dead030

25. Krawetz SA, Kruger A, Lalancette C, Tagett R, Anton E, Draghici S, et al. A survey of small RNAs in human sperm. Hum Reprod. 2011;26:3401–12. doi:10.1093/humrep/der329

26. Yeh LY, Lee RK, Lin MH, Huang CH, Li SH. Correlation between Sperm Micro Ribonucleic Acid-34b and -34c Levels and Clinical Outcomes of Intracytoplasmic Sperm Injection in Men with Male Factor Infertility. Int J Mol Sci. 2022;23:12381. doi:10.3390/ijms232012381

27. Yuan S, Tang C, Zhang Y, Wu J, Bao J, Zheng H, et al. mir-34b/c and mir-449a/b/c are required for spermatogenesis, but not for the first cleavage division in mice. Biol Open. 2015;4:212–23. doi:10.1242/bio.201410959

28. Wu J, Bao J, Kim M, Yuan S, Tang C, Zheng H, et al. Two miRNA clusters, miR-34b/c and miR-449, are essential for normal brain development, motile ciliogenesis, and spermatogenesis. Proc Natl Acad Sci U S A. 2014;111:E2851–7. doi:10.1073/pnas.1407777111

29. Dorostghoal M, Galehdari H, Hemadi M, Davoodi E. Correction to: Sperm miR-34c-5p Transcript Content and Its Association with Sperm Parameters in Unexplained Infertile Men. Reprod Sci. 2022;29:91. doi:10.1007/s43032-021-00770-5

30. Zhang W, Zhang Y, Zhao M, Ding N, Yan L, Chen J, et al. MicroRNA expression profiles in the seminal plasma of nonobstructive azoospermia patients with different histopathologic patterns. Fertil Steril. 2021;115:1197–211. doi:10.1016/j.fertnstert.2020.11.020

31. Shi S, Shi Q, Sun Y. The effect of sperm miR-34c on human embryonic development kinetics and clinical outcomes. Life Sci. 2020;256:117895. doi:10.1016/j.lfs.2020.117895

32. Zhang J, Liu X, Gao Y. The long noncoding RNA MEG3 regulates Ras-MAPK pathway through RASA1 in trophoblast and is associated with unexplained recurrent spontaneous abortion. Mol Med. 2021;27:70. doi:10.1186/s10020-021-00337-9

33. Chen S, Liu B, Li J, Liao S, Bi Y, Huang W, et al. Talin1 regulates endometrial adhesive capacity through the Ras signaling pathway. Life Sci. 2021;274:119332. doi: 10.1016/j.lfs.2021.119332

